## Supplemental Figure for "Comparative analysis of two *Caenorhabditis elegans* kinesins KLP-6 and UNC-104 reveals a common and distinct activation mechanism in kinesin-3"

### Supplementary figure S1

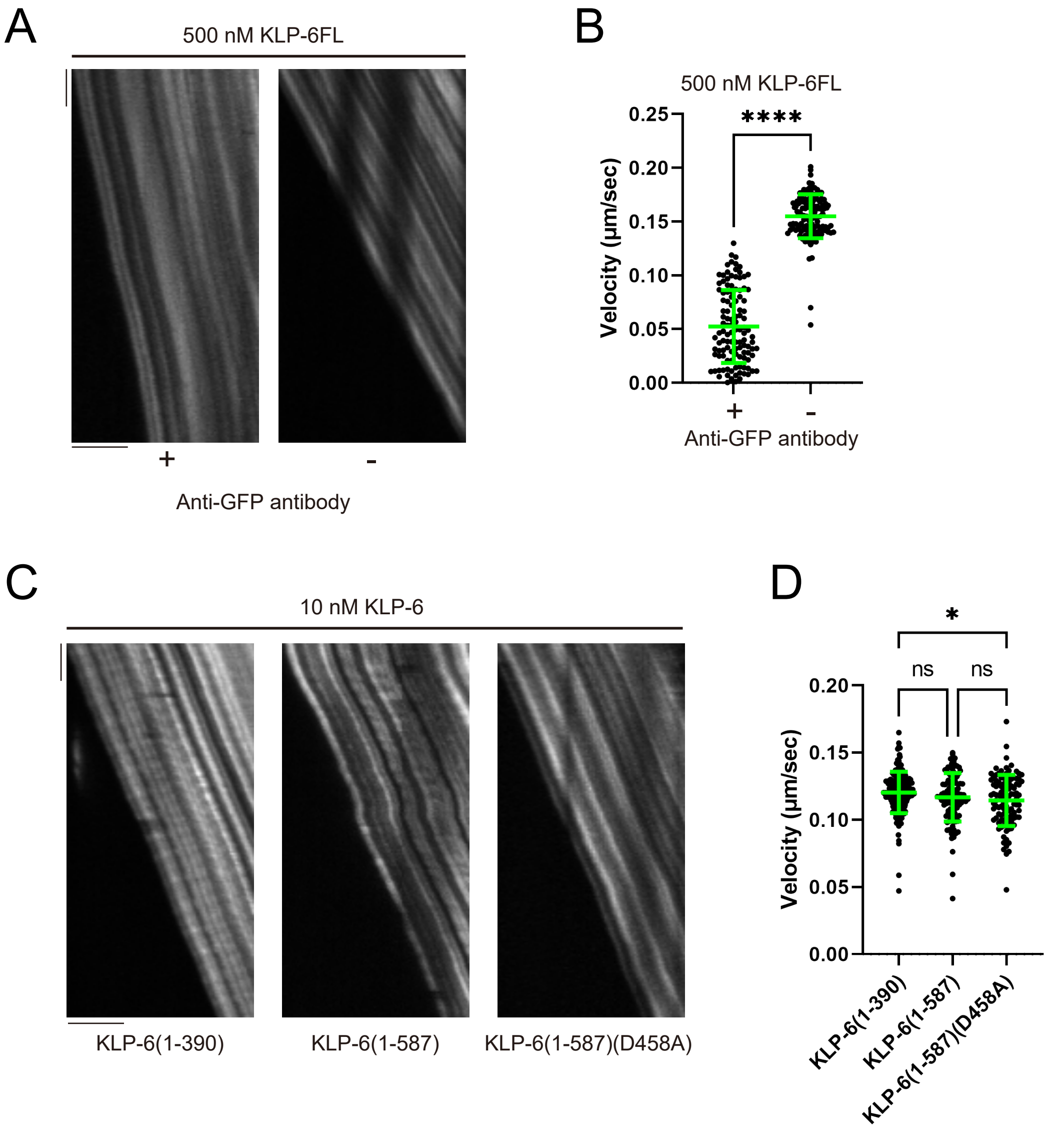

Supplementary Figure S2

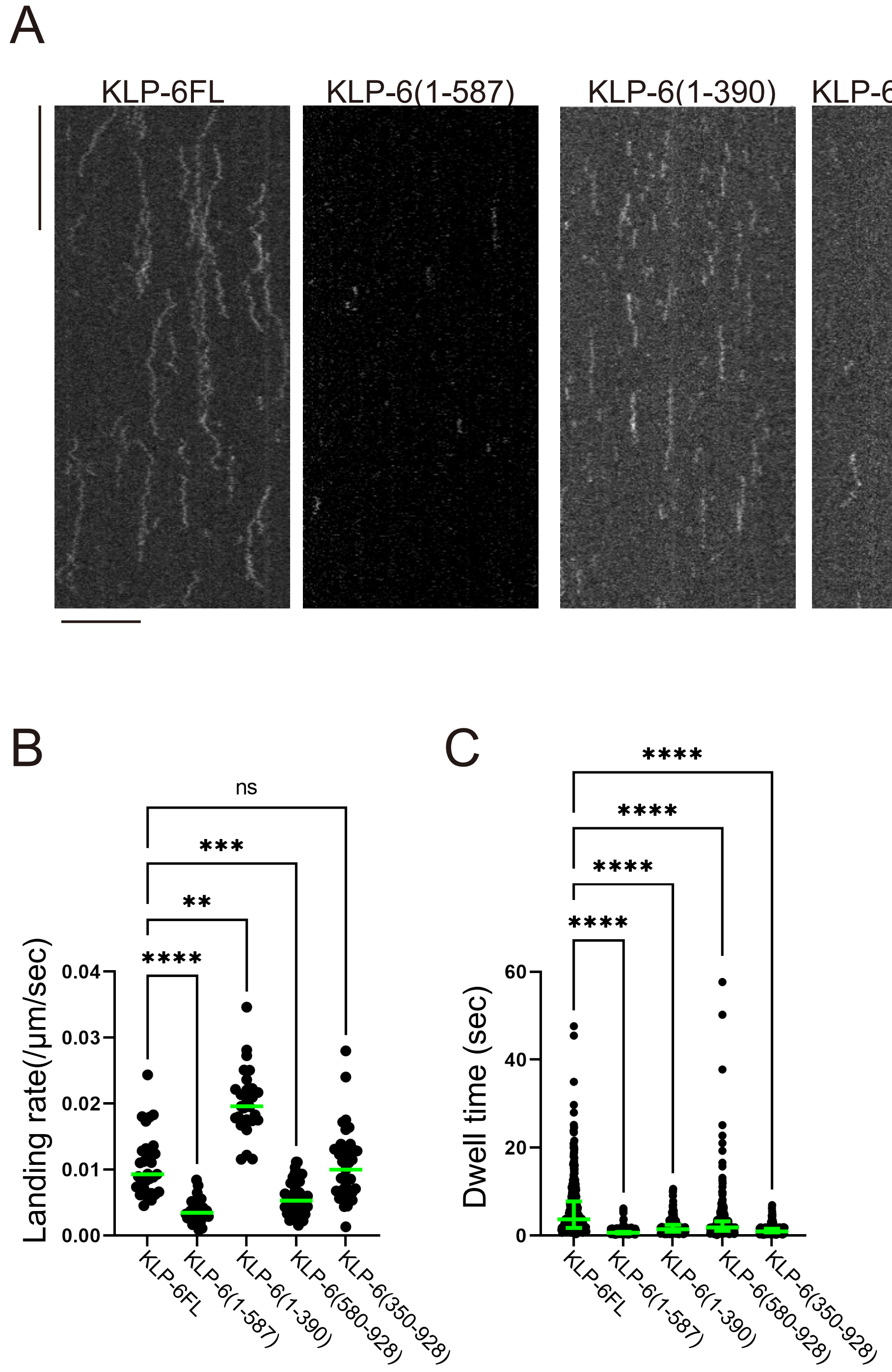

Supplementary figure S3

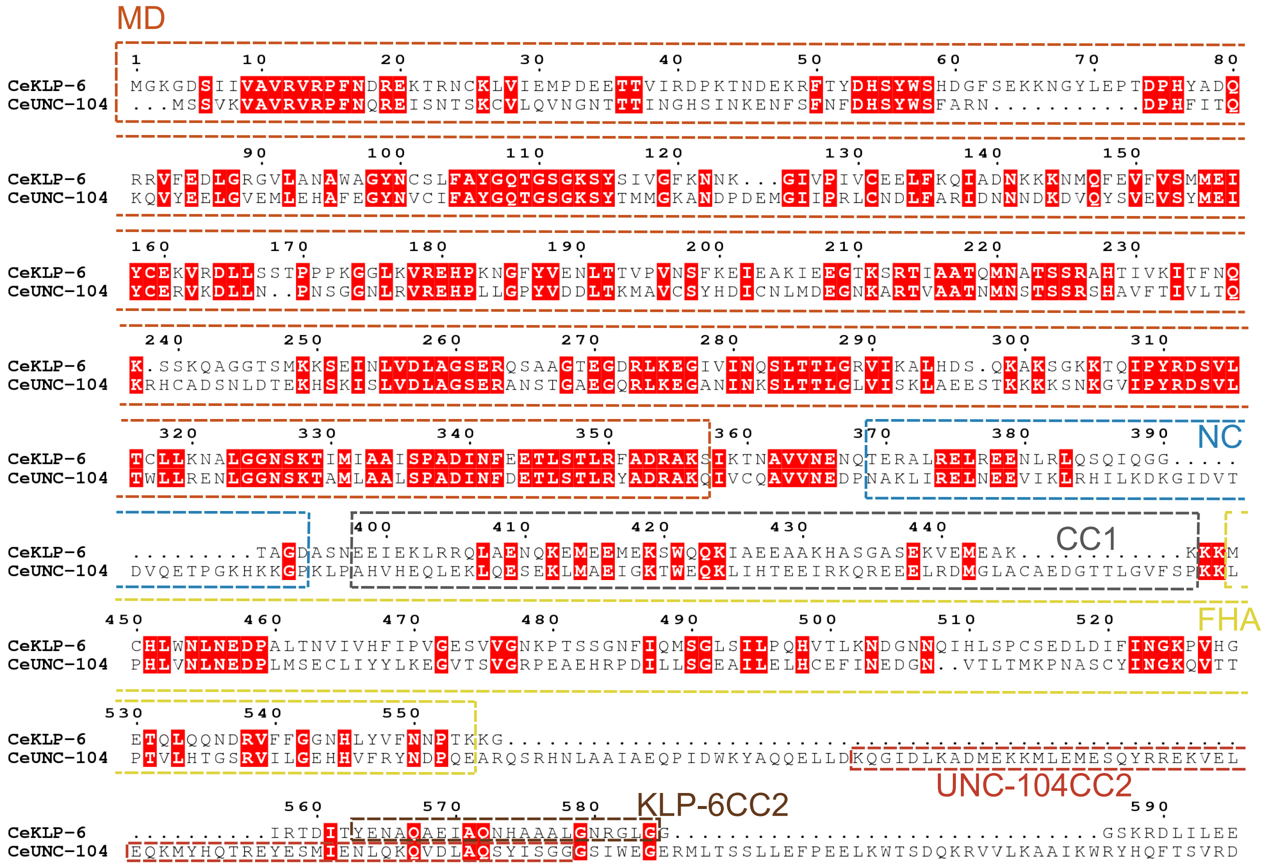

Supplementary figure S4

A

| | | | | |
|---|---|---|---|---|
| Ce UNC-104 | 407 - | LEKLQESEKLM | - | 417 |
| Ce KLP-6 | 404 - | RRQLAENQKEM | - | 414 |
| Hs KIF13B | 394 - | KDRLEESEKLI | - | 404 |

B

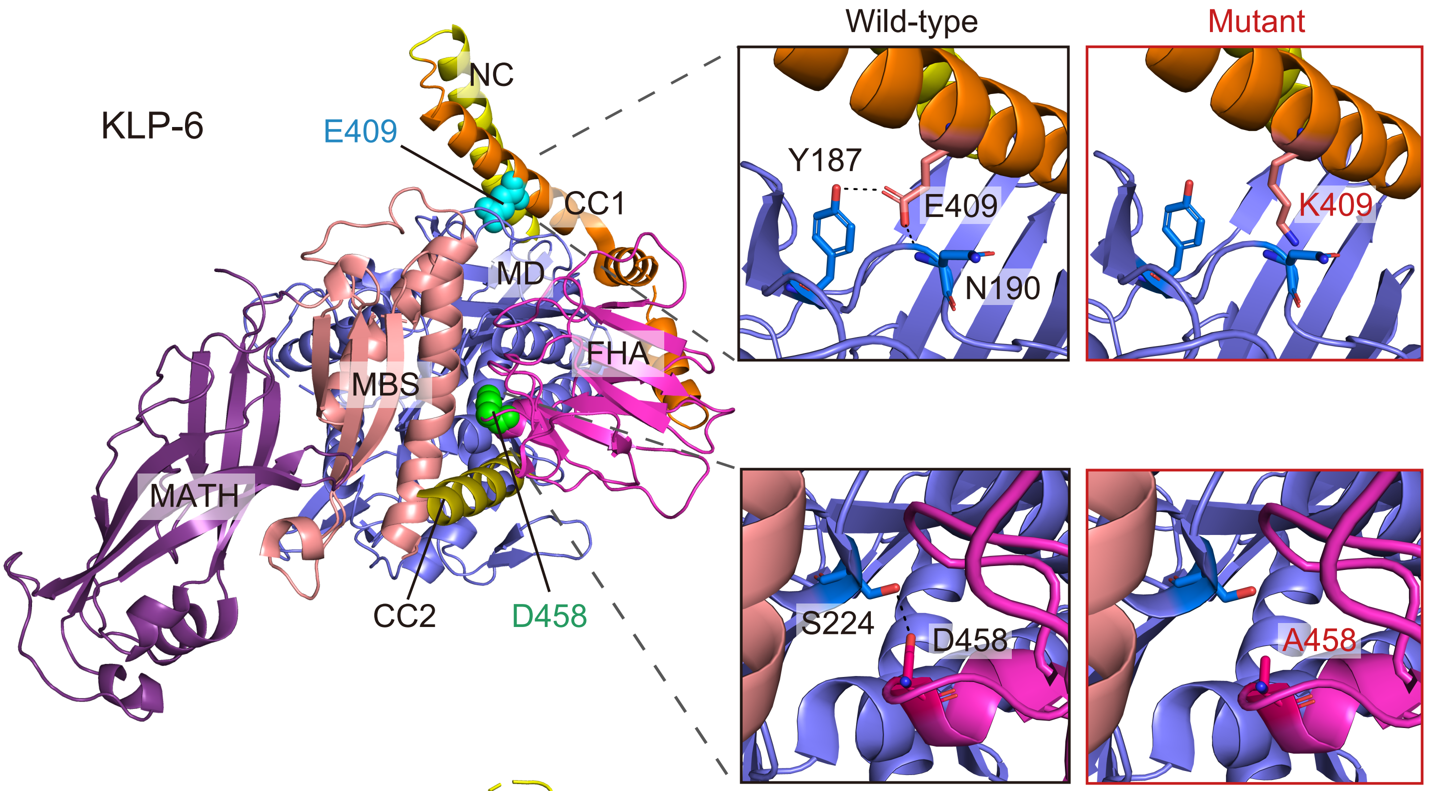


C

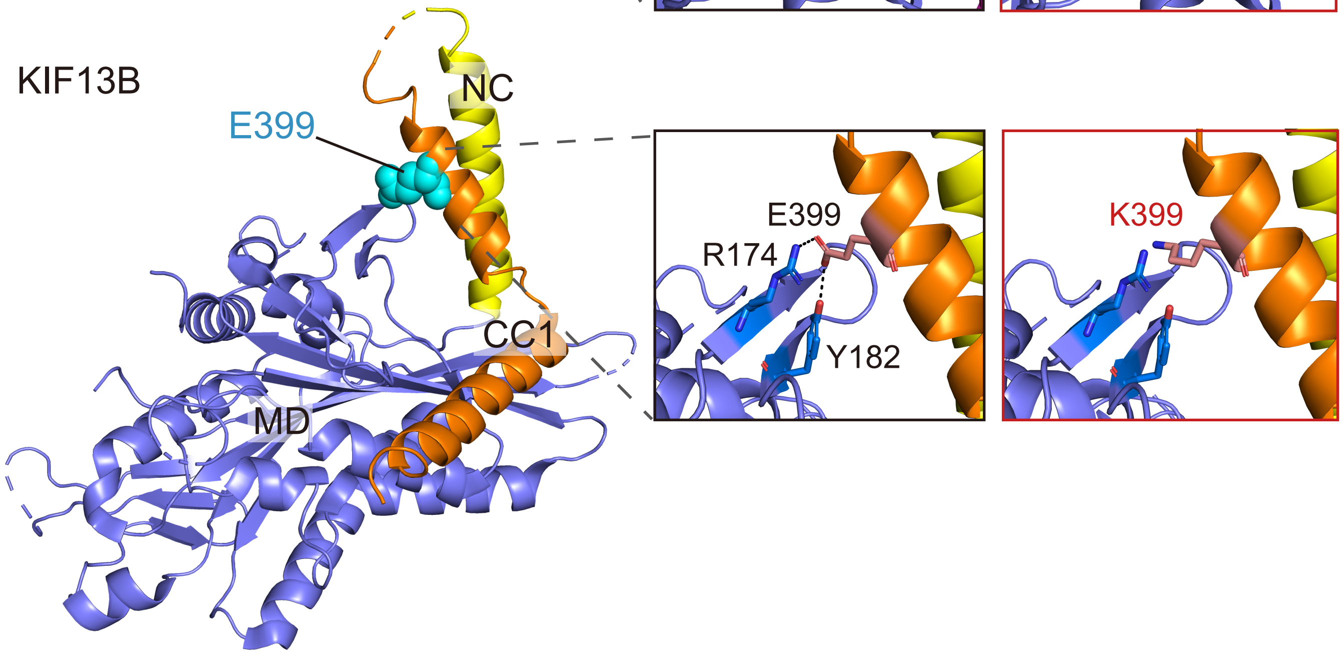

### Supplementary figure S5

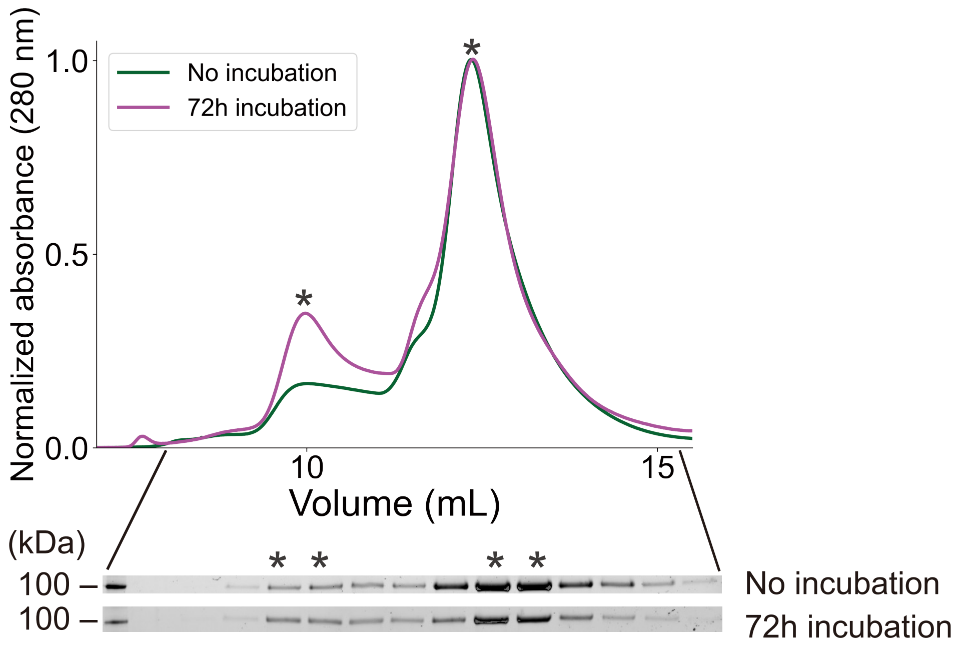

### Supplementary figure S6

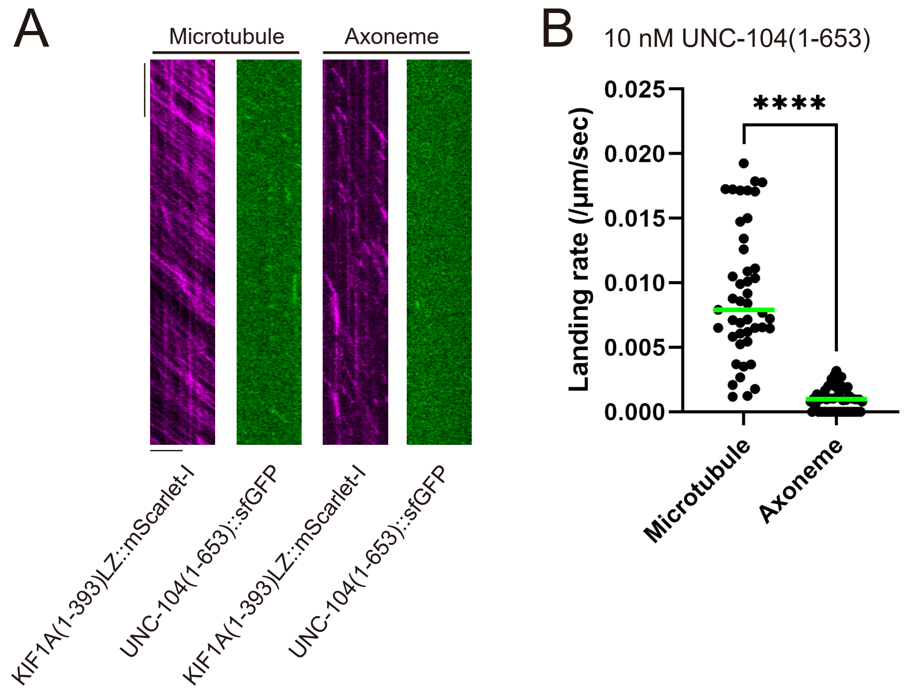

### Supplementary figure S7

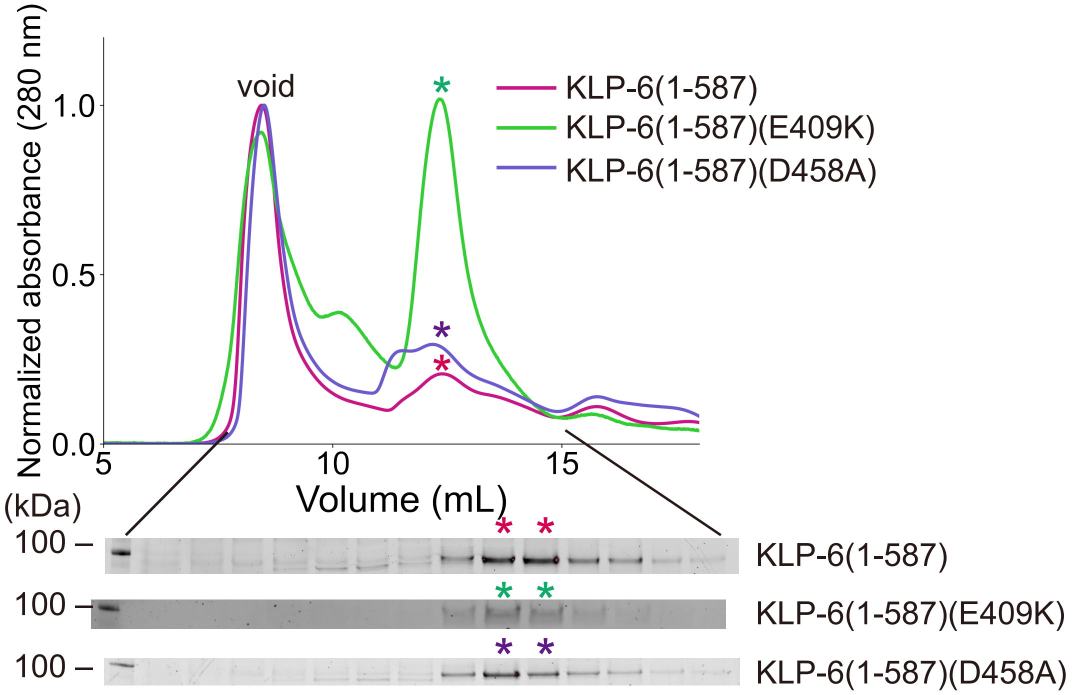

### Supplementary figure S8

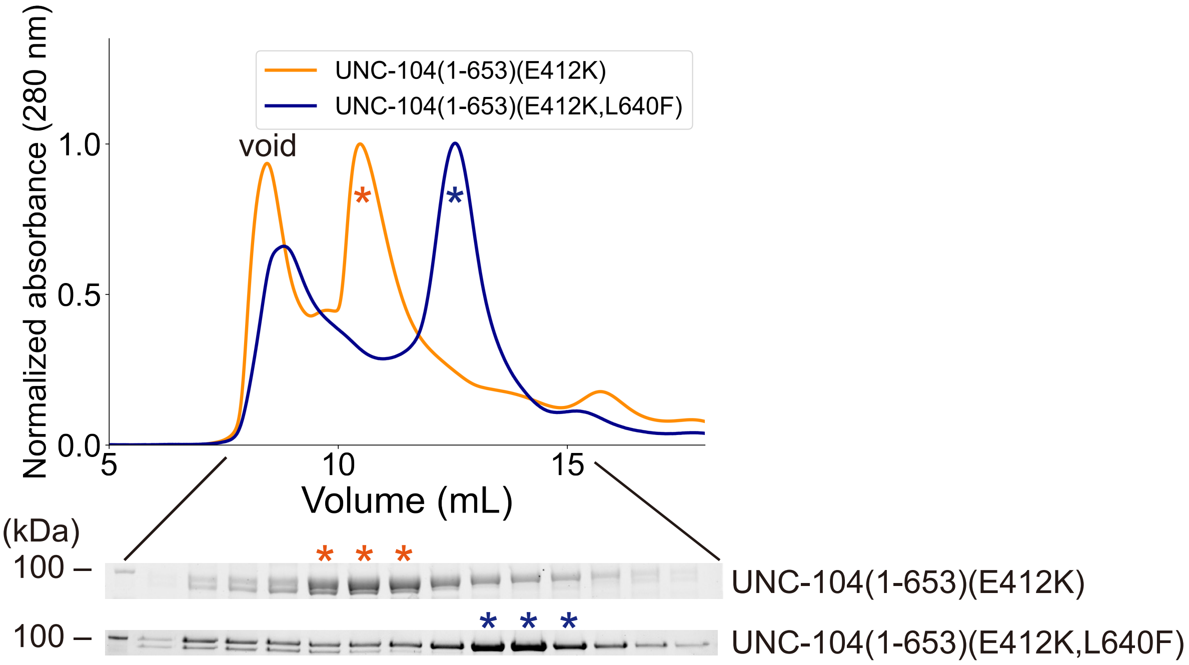

Supplementary figure S9

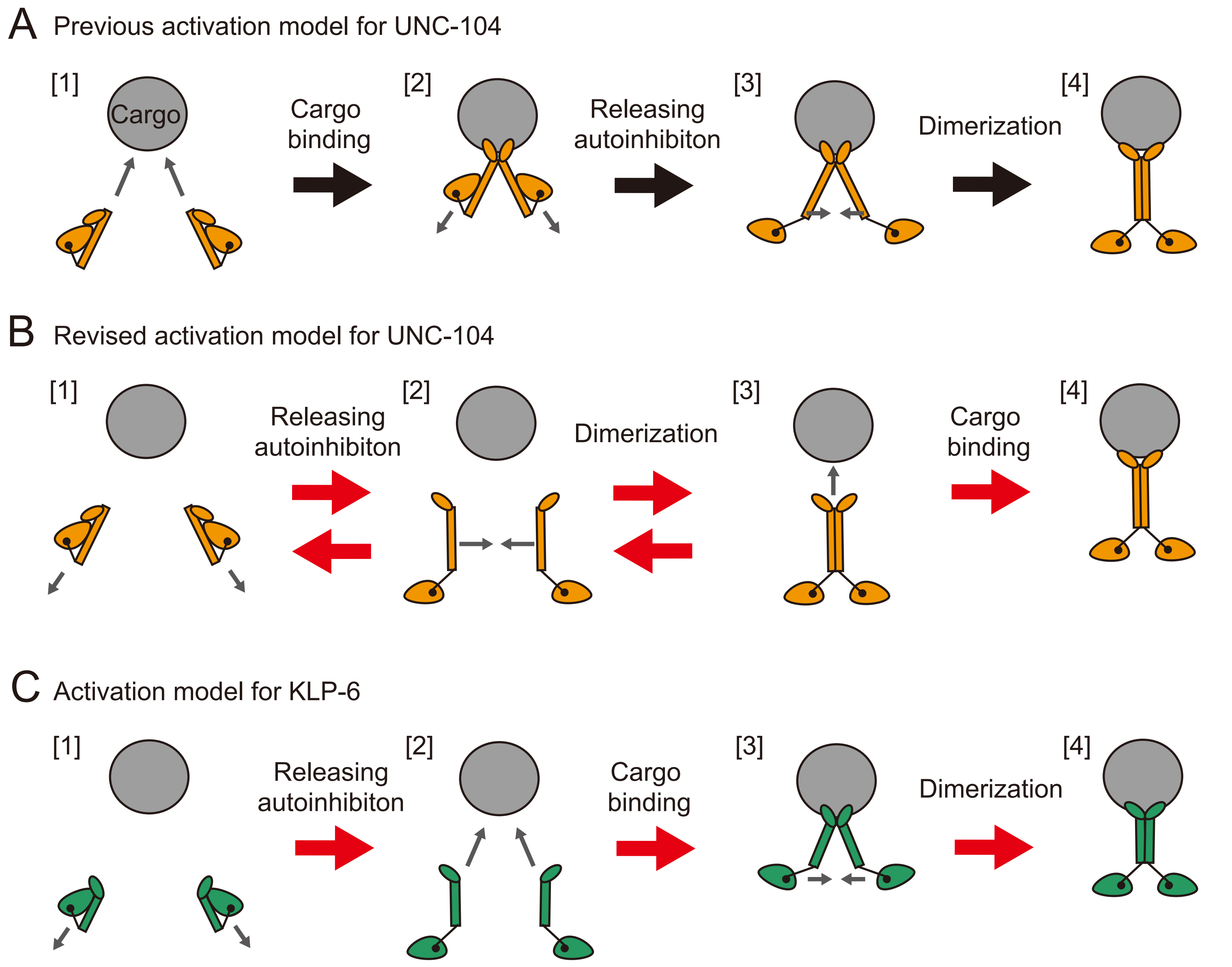

Supplementary figure S10

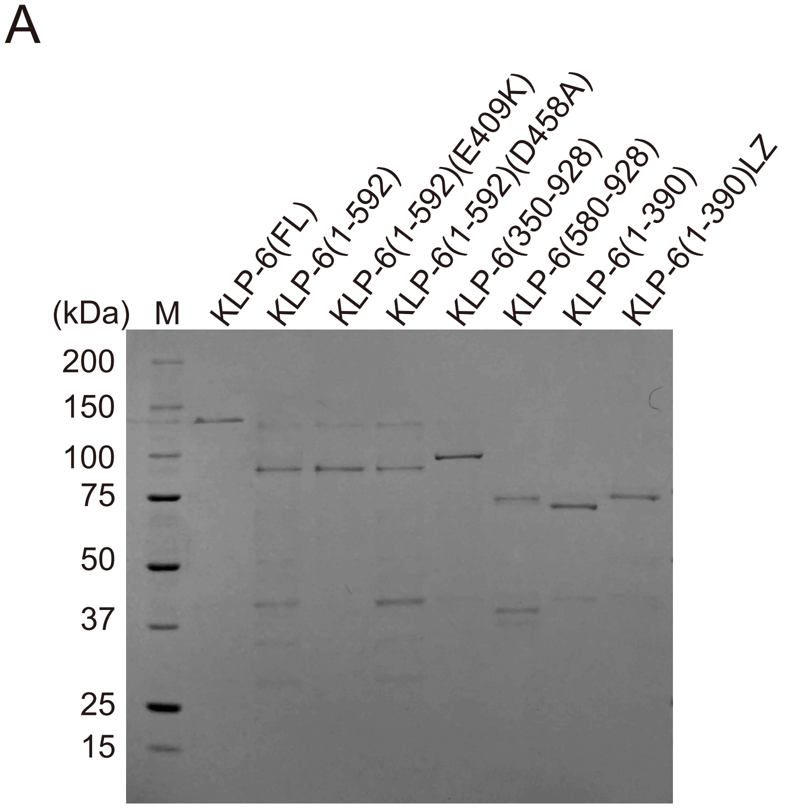


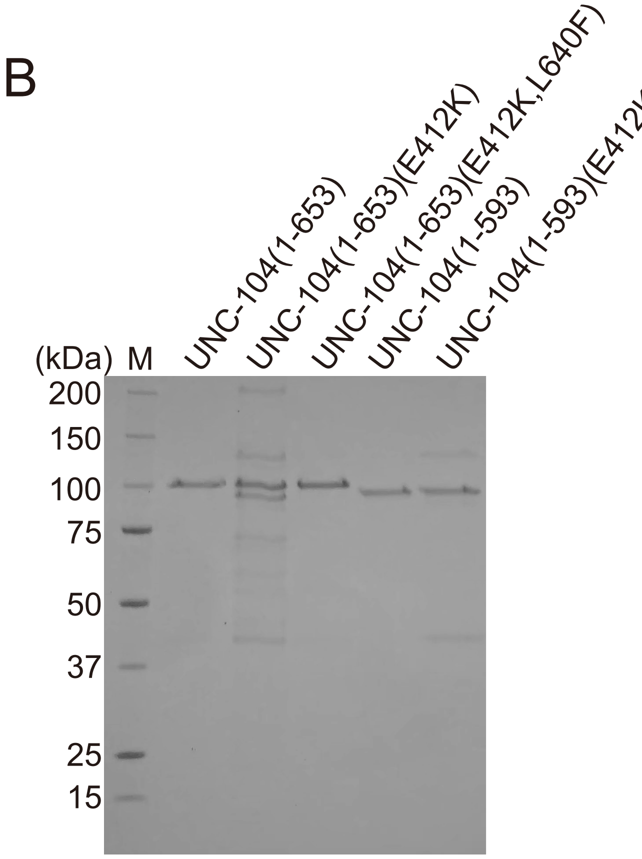
