## Supplemental Figure legends for "Comparative analysis of two *Caenorhabditis elegans* kinesins KLP-6 and UNC-104 reveals a common and distinct activation mechanism in kinesin-3"

**Supplementary Figure legends**

**Figure S1 Microtubule gliding assays using KLP-6.**

(A) Representative kymographs showing the microtubule gliding with 500 nM KLP-6FL. KLP-6FL was attached to the glass surface using anti-GFP antibody (+) or directly (-). Horizontal and vertical bars show 5 µm and 10 seconds, respectively.

(B) Dot plots showing the griding velocity of KLP-6FL when attached to the glass surface with (+) or without (-) anti-GFP antibody. Green bars represent mean ± S.D.. n = 115 and 127 microtubules for KLP-6FL with and without anti-GFP antibody. Student t-test. ****, p < 0.0001.

(C) Representative kymographs showing the microtubule gliding with 10 nM KLP-6(1-390), KLP-6(1-587) and KLP-6(1-587)(D458A). The motor was attached to the glass surface using anti-GFP antibody. Horizontal and vertical bars show 5 µm and 10 seconds, respectively. (D) Dot plots showing the griding velocity of 10 nM KLP-6(1-390), KLP-6(1-587) and KLP-6(1-587)(D458A). Each dot shows a single datum point. Each dot shows a single datum point. Green bars represent mean ± S.D.. n = 180, 107 and 103 microtubules for KLP-6(1-390), KLP-6(1-587) and KLP-6(1-587)(D458A). One-way ANOVA test followed by Tukey's multiple comparisons test. *, p < 0.05. ns, p > 0.05 and statistically not significant.

**Figure S2 Purified KLP-6 and its deletion mutants do not show processive runs**

(A) Representative kymographs showing the motility of 5 pM KLP-6FL, KLP-6(1-587), KLP-6(1-390), KLP-6(350-928) and KLP-6(580-928) on microtubules in the presence of 2 mM ATP. Horizontal and vertical bars show 10 µm and 10 seconds, respectively. None of the KLP-6 proteins shows directional movements on microtubules.

(B) Dot plots showing the landing rate of KLP-6FL, KLP-6(1-587), KLP-6(1-390), KLP-6(350-928) and KLP-6(580-928). Each dot shows a single datum point. Green bars represent median value. n = 31, 33, 27, 42 and 39 microtubules for KLP-6FL, KLP-6(1-587), KLP-6(1-390), KLP-6(350-928) and KLP-6(580-928). Kruskal-Wallis test followed by Dunn's multiple comparison test. **, p < 0.01, P < 0.001, and ****, p < 0.0001.

(C) Dot plots showing the dwell time on microtubules of KLP-6FL, KLP-6(1-587), KLP-6(1-390), KLP-6(350-928) and KLP-6(580-928). Each dot shows a single datum point. Green bars represent median value and interquartile range. n = 348 and 338, 326, 412 and 403 particles in KLP-6FL, KLP-6(1-587), KLP-6(1-390), KLP-6(350-928) and KLP-6(580-928), respectively. Kruskal-Wallis test followed by Dunn's multiple comparison test. ****, p < 0.0001.

**Figure S3 Comparison of KLP-6 and UNC-104**

Amino acid sequences of KLP-6 and UNC-104 are aligned using CLUSTALW on ESPript 3 (https://espript.ibcp.fr/ESPript/cgi-bin/ESPript.cgi). The motor domain, neck coil domain, CC1 domain, FHA domain and CC2 domains are indicated.

**Figure S4 Structural model of KLP-6 and KIF13B**

(A) Sequence comparison between Ce (*C. elegans*) UNC-104, Ce (*C. elegans*) KLP-6 and Hs (*Homo sapiens*) KIF13B. Ce UNC-104(E412) is conserved in Ce KLP-6(E409) and Hs KIF13B(E399). (B) Crystal structure of full-length KLP-6 (PDB 7WRG). The cyan sphere represents E409 from the CC1 domain, which interacts with Y187 and N190 from the motor domain (MD), while the mutation K409 does not interact with those residues. This mutation disrupts the MD-CC1 domain interaction and autoinhibition of KLP-6. The green sphere represents D458 from the FHA domain, which interacts with S224 from the MD, while the mutation A458 does not interact with this residue. This mutation disrupts the MD-FHA domain interaction and autoinhibition of KLP-6. (C) Crystal structure of MD-NC-CC1 of KIF13B (PDB 6A20). The cyan sphere represents E399 from the CC1 domain, which interacts with R174 and Y182 from the MD, while the mutation K399 does not interact with those residues. This mutation disrupts the MD-CC1 domain interaction and autoinhibition of KIF13B. The mutations were generated using PyMOL.

**Figure S5 Monomer-dimer re-equilibrium of monomeric UNC-104(1-653)**

Size exclusion chromatography of the monomer fraction of UNC-104(1-653), indicated by the blue asterisk in Fig. 5B in the main text. Without incubation (green), the dimer peak (left asterisk) was minimal, but after 72 hours of incubation at 4 ℃ (plum), the peak increased. The SDS-PAGE of the elution fractions are shown beneath the profile. The number shown at the left side indicates molecular weight standard.

**Figure S6 Single-molecule analysis using *Chlamydomonas* axonemes**

(A) Representative kymographs showing the motility of 0.2 nM KIF1A(1-393)LZ::mScarlet-I and 10 nM UNC-104(1-653)::sfGFP on purified porcine microtubules within the same chamber, as well as the motility of 0.2 nM KIF1A(1-393)LZ::mScarlet-I and 10 nM UNC-104(1-653)::sfGFP on *Chlamydomonas* axonemes within the same chamber. Horizontal and vertical bars show 5 µm and 5 seconds, respectively. UNC-104(1-653)::sfGFP shows no processive runs on *Chlamydomonas* axonemes. (D) Dot plots showing the landing rate of UNC-104(1-653)::sfGFP on purified porcine microtubules and *Chlamydomonas* axonemes. Each dot shows a single datum point. Green bars represent median value. n = 45 and 46 microtubules for UNC-104(1-653)::sfGFP on purified porcine microtubules and *Chlamydomonas* axonemes, respectively. Mann-Whitney U test. ****, p < 0.0001.

**Figure S7 KLP-6(E409K) does not form a dimer**

Size exclusion chromatography of KLP-6(1-587) (plum), KLP-6(1-587)(E409K) (green) and KLP- 6(1-587)(D458A) (purple). The SDS-PAGE of the elution fractions are shown beneath the profile. Data for KLP-6(1-587) and KLP-6(1-587)(D458A) are replotted and reshown from Figure 4B. Note that no significant differences are observed among the elution profiles of the three proteins. The number shown at the left side indicates molecular weight standard.

**Figure S8 UNC-104(E412K,L640F) does not form a dimer**

Size exclusion chromatography of UNC-104(1-653)(E412K) (orange) and UNC-104(1-653)(E412K,L640F) (navy blue). The SDS-PAGE of the elution fractions are shown beneath the profile. Data for UNC-104(1-653)(E412K) are replotted and reshown from Figure 5B. Note that UNC-104(1-653)(E412K,L640F) predominantly eluted in the monomer peak, in contrast to UNC-104(1-653)(E412K), which predominantly eluted in the dimer peak. The number shown at the left side indicates molecular weight standard.

**Figure S9** Activation models for UNC-104 and KLP-6

(A) Previous activation model for UNC-104. UNC-104 initially exists in a monomeric state in solution and subsequently binds to cargo (transition 1 → 2), triggering the release of its autoinhibition (transition 2 → 3). On the cargo membrane, the motor can then undergo dimerization (transition 3 → 4), initiating processive transport on microtubules.

(B) Proposed activation model for UNC-104. Upon release of autoinhibition (transition 1 → 2), UNC-104 can dimerize independently of cargo binding (transition 2 → 3). Note that the transitions 1 → 2 and 2 → 3 are reversible. Following dimerization, the motor can bind to cargo and initiate processive transport on microtubules.

(C) Proposed activation model for KLP-6. KLP-6 is strongly autoinhibited and primarily exists as a monomer in solution. Upon release of autoinhibition (transition 1 → 2), KLP-6 engages with cargo (transition 2 → 3), facilitating subsequent dimerization of the motor (transition 3 → 4). Subsequently, the motor can initiate processive transport on microtubules.

**Figure S10 Proteins analyzed in this study**

Full scan images of SDS-PAGE showing the purified proteins analyzed in this study. Lane M, molecular weight markers. Numbers shown at the left side indicate the molecular weight standard.

(A) KLP-6 proteins.

(B) UNC-104 proteins.
